# Estimating the true stability of the prehydrolytic outward-facing state in an ABC protein

**DOI:** 10.1101/2023.07.16.549217

**Authors:** Márton A. Simon, Iordan Iordanov, András Szöllősi, László Csanády

## Abstract

CFTR, the anion channel mutated in cystic fibrosis patients, is a model ABC protein whose ATP-driven conformational cycle is observable at single-molecule level in patch-clamp recordings. Bursts of CFTR pore openings are coupled to tight dimerization of its two nucleotide binding domains (NBDs) and in wild-type (WT) channels are mostly terminated by ATP hydrolysis. The slow rate of non-hydrolytic closure – which determines how tightly bursts and ATP hydrolysis are coupled – is unknown, as burst durations of catalytic site mutants span a range of ∼200-fold. Here we show that Walker A mutation K1250A, Walker B mutation D1370N, and catalytic glutamate mutations E1371S and E1371Q all completely disrupt ATP hydrolysis. True non-hydrolytic closing rate of WT CFTR approximates that of K1250A and E1371S. That rate is slowed ∼15-fold in E1371Q by a non-native inter-NBD H-bond, and accelerated ∼15-fold in D1370N. These findings uncover unique features of the NBD interface in human CFTR.

## Introduction

The CFTR anion channel is a key component of transepithelial salt-water transport in the lung, the pancreas and the intestine, and its mutations cause cystic fibrosis (O’Sullivan and Freedman, 2009). CFTR (ABCC7) is the only known ion channel member of the family of ATP Binding Cassette (ABC) proteins. Its two ABC-typical halfs, each consisting of a transmembrane domain (TMD1 and 2, Fig. 1A, *gray*) followed by a cytosolic nucleotide binding domain (NBD1 and 2; Fig. 1A, *blue* and *green*), are linked by a unique regulatory (R) region (Fig. 1A, *pale red*) which must be phosphorylated by cAMP-dependent protein kinase (PKA) to allow channel activity (Riordan et al., 1989; Cheng et al., 1991).

**Fig. 1.**
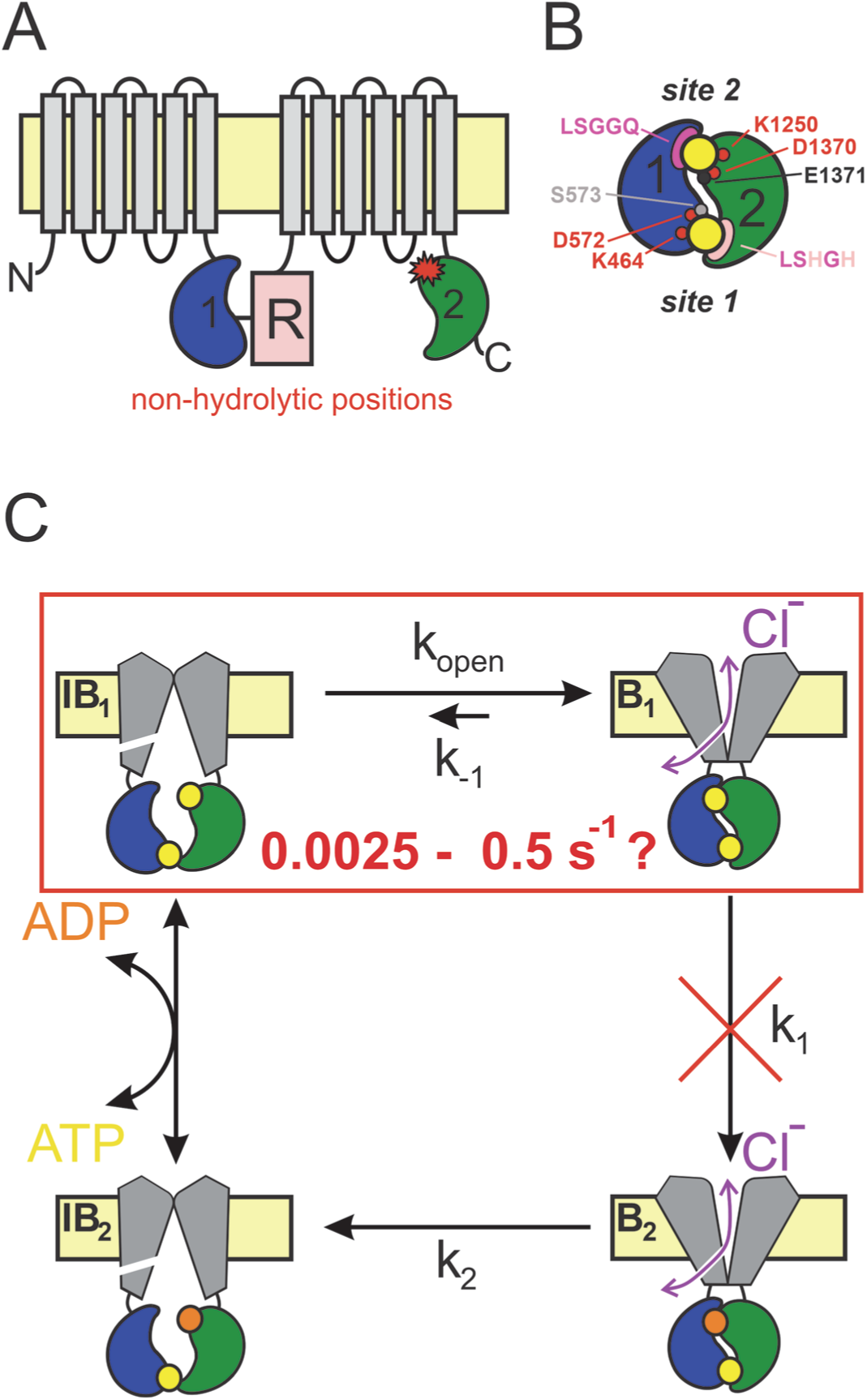
CFTR topology and gating cycle. *A*, Cartoon topology of CFTR. TMD1-2, *gray*; NBD1, *blue*; NBD2, *green*; R-domain, *pale red*. *Red asterisk* denotes catalytic site mutations. *B*, Cartoon of head-to-tail NBD dimer. Color coding as in *A*. ATP, large yellow circles. Site 2 (*top*): conserved Walker A/B residues, *small red circles*; catalytic base, *small black circle*; signature sequence, *magenta crescent*. Site 1 (*bottom*): color coding as for site 2, *pale residues* represent non-conserved substitutions. *C*, Schematic gating cycle of phosphorylated CFTR. Flickery closures from states B_1_ and B_2_ are not depicted. Disruption of ATP hydrolysis in catalytic site mutants (*red cross*) reduces gating to reversible IB_1_↔B_1_ transitions (*red box*). Note, that in reality sites 1 and 2 are equally near the membrane, in the cartoons site 2 is depicted as the top and site 1 as the bottom site merely for representation purposes.

ABC proteins are present in all kingdoms of life. The 48 human ABC proteins play essential physiological roles by mediating transmembrane transport of a variety of substrates (Dean et al., 2022). Within the ABC family the NBDs are the most highly conserved modules, and consist of a core subdomain that binds ATP (the "head") and an ABC-specific alpha-helical subdomain (the "tail"). In all ABC proteins in the presence of ATP the two NBDs form a tight head-to-tail dimer which occludes two ATP molecules (Fig. 1B, *yellow*) at the interface. Both nucleotide binding sites are flanked on one side by the conserved Walker A and Walker B motifs in the head of one NBD (Fig. 1B, *red*), and on the other side by the conserved "ABC signature sequence" in the tail of the other NBD (Fig. 1B, *magenta*). The ATP-bound NBD dimer is stable, but is disrupted following ATP hydrolysis (Locher, 2016). In a subset of ABC proteins, which includes the entire C subfamily, only one of these two composite ATP binding sites is catalytically active (Procko et al., 2009). In CFTR the composite site formed by the head of NBD2 and the tail of NBD1 ("site 2", Fig. 1B, *upper site*) contains canonical sequence motifs and hydrolyses ATP with an overall turnover number of ∼0.5-1 s^-1^ (Li et al., 1996; Liu et al., 2017). In contrast, the other site ("site 1", Fig. 1B, *lower site*) has accumulated non-canonical substitutions in several key motifs (Fig. 1B, *pale residues*) and is catalytically inactive (Aleksandrov et al., 2002; Basso et al., 2003).

During substrate transport the TMDs of ABC exporters alternate between inward-facing (IF) and outward-facing (OF) conformations. In a phosphorylated CFTR channel opening and closure of the transmembrane ion pore ("gating") follows analogous TMD movements (Csanády et al., 2019) (Fig. 1C). The IF conformation, in which the pore is sealed near its extracellular end (Fig. 1C, *left*), is a long-lived state (∼1 s) and corresponds to long "interburst" (IB) closed dwell times observable in single-channel current recordings. In the OF conformation the external gate is predominantly open, and a lateral portal which connects the channel pore with the cytosol generates a transmembrane aqueous pathway permeable to anions ((Zhang et al., 2018); Fig. 1C, *right*, *double arrows*). The OF state is also relatively stable (dwell time hundreds of milliseconds), but the continuity of its transmembrane pore is occasionally disrupted for brief (∼10 ms) intervals by a smaller conformational change (not depicted in Fig. 1C) likely confined to the external ends of the TMD helices (Zhang et al., 2017; Zhang et al., 2018; Simon and Csanády, 2021). Correspondingly, in single-channel recordings the OF state corresponds to "bursts" (B) of channel openings separated by brief ("flickery") closures (Winter et al., 1994).

In ABC exporters NBD and TMD movements are coupled: formation of the tight ATP-bound NBD dimer flips the TMDs from an inward-facing (IF) to an outward-facing (OF) conformation, and disruption of the tight NBD dimer following ATP-hydrolysis resets the TMDs to the IF conformation (Locher, 2016). Gating of phosphorylated CFTR channels is driven by an analogous unidirectional conformational cycle (Csanády et al., 2019) (Fig. 1C). In all ABC-C proteins the OF state of the TMDs is coupled to tight dimerization of both NBD catalytic sites (Johnson and Chen, 2018; Zhang et al., 2018; Huang et al., 2023). Correspondingly, functional studies on CFTR showed that site 2 is tightly dimerized in the B state (Vergani et al., 2005) (Fig. 1C, *right*). In contrast, in IF structures of ABC-C proteins obtained in the absence of ATP (Johnson and Chen, 2017; Liu et al., 2017; Huang et al., 2023), or for unphosphorylated CFTR even in its presence (Levring et al., 2023), the NBDs are seen to widely separate losing all contact across both composite sites. Interestingly, in the asymmetric bacterial ABC protein Tm287-288 some contacts across the degenerate site are retained throughout the entire conformational cycle, even in the IF state, as shown by extensive structural (Hohl et al., 2012; Hohl et al., 2014) and functional (Timachi et al., 2017) studies. Similarly, for phosphorylated CFTR channels functional studies suggested that site 1 does not completely separate throughout the entire ATP-driven gating cycle, even during IB events (Tsai et al., 2010; Szollosi et al., 2011) (see NBDs in Fig. 1C, *left*), a prediction confirmed by recent single-molecule FRET measurements (Levring et al., 2023).

In ABC exporters thermodynamically uphill transport requires unidirectional conformational cycling. The "coupling ratio" (CR), i.e., the fraction of initiated cyles that are completed through ATP hydrolysis, depends on the relative rates of the two possible exit pathways from the prehydrolytic OF state: ATP hydrolysis (rate *k*_1_, Fig. 1C) vs. non-hydrolytic NBD dimer dissociation (*k*_-1_, Fig. 1C). Because typically *k*_1_>>*k*_-1_, CR=*k*_1_/(*k*_1_+*k*_-1_) is near unity. In most ABC transporters the actual values of these rates are hard to directly estimate, but in CFTR direct measurements of conformational dwell-times are made feasible by single-channel current recordings. For a wild-type (WT) channel the mean burst duration (τ_b_) reports the sum of the life times of the prehydrolytic (Fig. 1C, state B_1_) and posthydrolytic (Fig. 1C, state B_2_) OF conformations. Because the life time of B_2_ is short compared to that of B_1_ (*k*_2_>>*k*_1_; Fig. 1C; (Gunderson and Kopito, 1995; Csanády et al., 2010)), 1/τ_b_ provides a rough estimate of rate *k*_1_, which is ∼4-5 s^-1^ at room temperature for pre-phosphorylated CFTR channels gating in ATP. In contrast, rate *k*_-1_ in WT CFTR is difficult to directly assess, and its estimates are based on the burst-state stability of various catalytic site mutants (Fig. 1A, *red star*) in which ATP hydrolysis is disrupted (Fig. 1C, *red cross*) and gating in saturating ATP is thus reduced to reversible IB_1_↔B_1_ transitions (Fig. 1B, *red box*). Numerous studies in the past have employed multiple different mutations for that purpose, including NBD2 Walker A lysine mutation K1250A (Gunderson and Kopito, 1995; Carson et al., 1995; Zeltwanger et al., 1999; Vergani et al., 2003; Csanády et al., 2006; Csanády et al., 2013; Csanády and Torocsik, 2014), Walker B aspartate mutation D1370N (Gunderson and Kopito, 1995; Bompadre et al., 2005; Csanády et al., 2010; Sorum et al., 2015; Yeh et al., 2015; Sorum et al., 2017; Simon and Csanády, 2021), and E1371S/Q mutations which eliminate the catalytic base (Vergani et al., 2003; Bompadre et al., 2005; Zhou et al., 2005; Vergani et al., 2005; Zhou et al., 2006; Csanády et al., 2013; Yu et al., 2016). Of note, so far all OF structures of CFTR (PDBID: 6msm, 6o2p, 6o1v, 7svd, 7sv7), as well as OF structures of several other ABC proteins (6bhu, 6cov, 6hbu, 6s7p, 7ekl, 8iza), were obtained from E-to-Q catalytic glutamate mutants.

However, interpretation of such mutant data is hampered by the fact that – even when compared under identical experimental conditions (expression system, temperature, phosphorylation state) – non-hydrolytic closing rates of the above mutants scatter over a range of ∼200-fold, from ∼0.0025 s^-1^ s for E1371Q (Vergani et al., 2005; Yu et al., 2016) to ∼0.5-1 s^-1^ for D1370N (Vergani et al., 2003; Yeh et al., 2015). The reason for that large scatter is unknown. One extreme possibility is that ATP hydrolysis is completely abolished only in the slowest-closing mutant (E1371Q) which therefore faithfully reports WT B_1_-state stability, whereas in the faster-closing mutants significant residual hydrolytic activity persists. Such a scenario would call into question conclusions based on the assumption of equilibrium gating in the latter mutants (Sorum et al., 2015; Yeh et al., 2015; Sorum et al., 2017; Simon and Csanády, 2021). The other extreme possibility is that all of the above mutations eliminate ATP hydrolysis, but different mutations differentially affect – either increase or decrease – B_1_-state stability. In either case, estimating the true stability of state B_1_ (i.e., rate *k*_-1_) for WT CFTR would substantially promote our understanding of gating energetics. The aim of the present study was to estimate the true rate *k*_-1_ for a pre-phosphorylated WT CFTR channel gating in ATP, as well as to verify disruption of ATP hydrolysis and evaluate effects on B_1_-state stability of the various non-hydrolytic mutations that have been extensively employed as models in the past. The results suggest unique features of the site-2 NBD interface in human CFTR compared to other ABCC family proteins.

## Results

### Interfacial H-bond between NBD1 D-loop and the mutated NBD2 catalytic glutamate side chain suggested by structures of outward-facing human, but not zebrafish, E-to-Q CFTR

The life time of the B_1_ state of non-hydrolytic CFTR mutants can be conveniently measured in inside-out macro-patch recordings following activation of pre-phosphorylated CFTR channels by a brief exposure to ATP: the time constant of the macroscopic current relaxation following ATP removal reports τ_b_. A previous study that compared NBD dimer stabilities of the human and zebrafish orthologs of CFTR (hCFTR and zCFTR) uncovered a puzzling discrepancy between the two CFTR variants. Whereas for hCFTR a large difference between the burst durations of the serine vs. glutamine mutant of the catalytic glutamate (hE1371S vs. hE1371Q) had long been documented (cf., Fig. 2A, *black* vs. *dark blue trace*; Fig. 2B, *black* vs. *dark blue symbols*), no significant difference (p=0.092) between τ_b_ of the analogous mutants of the zebrafish ortholog (zE1372S vs. zE1372Q; Fig. 2A, *gray* vs. *light blue trace*; Fig. 2B, *gray* vs. *light blue symbols*) was detectable (Simon and Csanády, 2023).

**Fig. 2.**
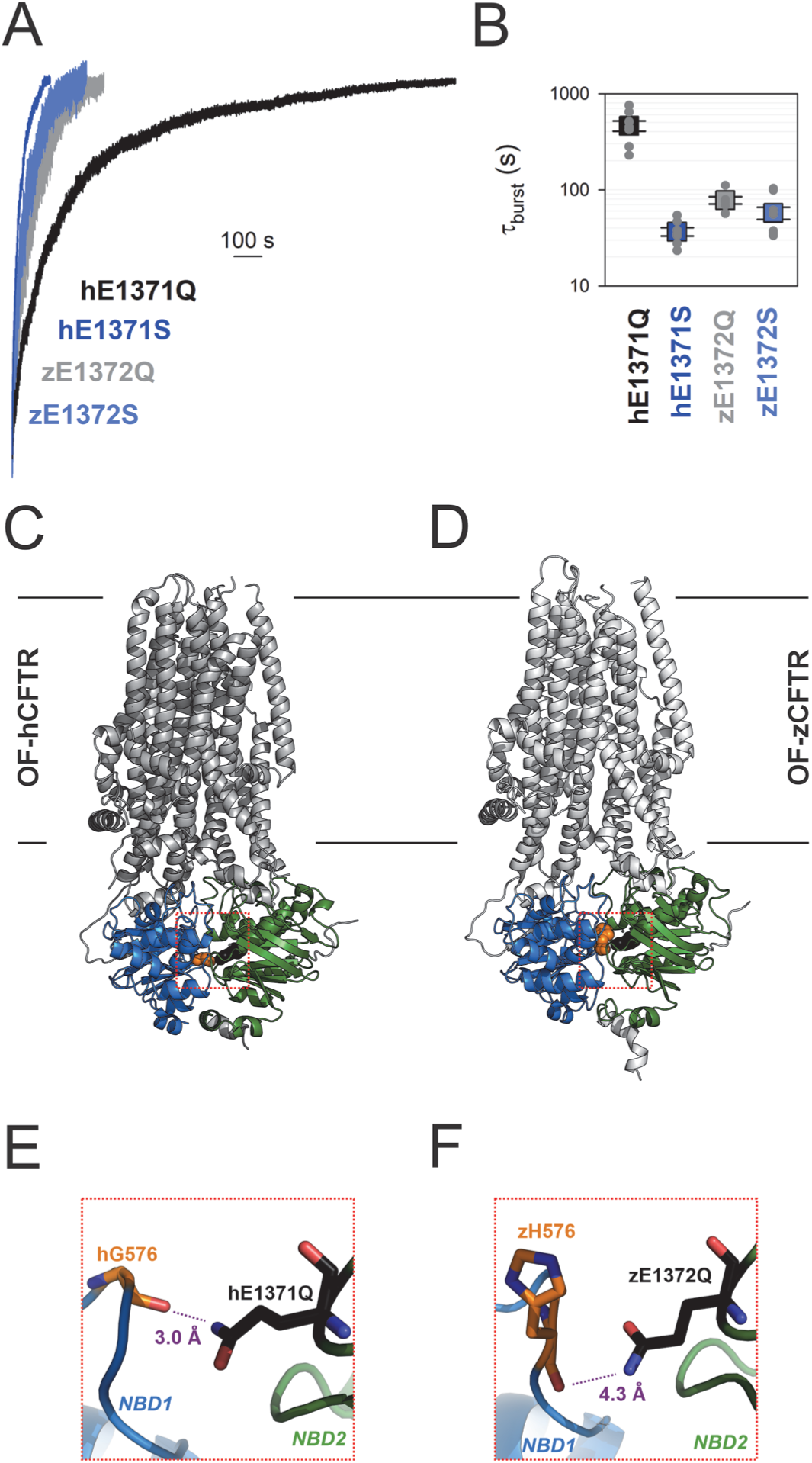
Non-native inter-NBD H-bond in OF E-to-Q mutant hCFTR, but not zCFTR, suggested by cryo-EM structures. *A*, Macroscopic current relaxations upon ATP removal in inside-out patches excised from *Xenopus laevis* oocytes expressing hE1371Q (*black trace*), hE1371S (*dark blue trace*), zE1372Q (*gray trace*), and zE1372S (*light blue trace*) CFTR channels. Currents were activated by exposure to 5 mM ATP, following prior phosphorylation for ∼2 minutes with 300 nM bovine PKA catalytic subunit. Current amplitudes are shown rescaled by their steady-state values, membrane potential (V_m_) was -20 to -80 mV. *B*, Mean burst durations (in seconds) obtained as the time constants of single-exponential fits to the macroscopic relaxations shown in *A*. Data are shown as mean±S.E.M. from 6-9 experiments using a logarithmic ordinate. Data in *A*-*B* are replotted from (Simon and Csanády, 2023). *C*-*D*, Cryo-EM structures of hCFTR-E1371Q (PDBID: 6msm) and zCFTR-E1372Q (PDBID: 5w81) in the OF conformation. Color coding as in *A*, positions hG576/zH576 (*orange*), and hQ1371/zQ1372 (*black*) are shown in spacefill. *Red dotted squares* identify regions expanded in panels *E*-*F*. *E*-*F*, Close-up views of the regions surrounding the mutated catalytic glutamate side chains (*black sticks*) in OF hCFTR (*E*) and zCFTR (*F*).

Those unexpected findings suggested differences in the fine structure of the site-2 interface in the bursting state of the two orthologs. We therefore examined the OF cyoelectronmicroscopy (cryo-EM) structures of the E-to-Q mutants of both orthologs (Fig. 2C-D) in more detail. In hCFTR (PDBID: 6msm) close examination identified a hydrogen bond across the site-2 interface, formed between the amide nitrogen of the engineered hQ1371 side chain and the peptide carbonyl oxygen of residue hG576, located in the opposing D-loop of NBD1 (Fig. 2E; cf., electron density in Fig. 2 – Fig. Suppl. 1). That H-bond cannot be present in WT CFTR, as the native hE1371 side chain contains no chemical moiety suitable to form a stabilizing interaction with the opposing carbonyl group. Interestingly, in the zCFTR structure (PDBID: 5w81) the distance between the mutant zQ1372 side chain and the nearest peptide carbonyl oxygen in the NBD1 D-loop is substantially larger (4.3A vs. 3.0 A), precluding formation of an H-bond (Fig. 2F). These observations raised the possibility that the >10-fold longer lifetime of the bursting state of hE1371Q compared to hE1371S CFTR (Fig. 2A-B) reflects artificial stabilization of state B_1_ by the introduced non-native interfacial H-bond.

### In E1371Q-hCFTR the Q1371 side-chain amino and G576 peptide carbonyl groups are energetically coupled in the bursting state

Gating-state dependent changes in energetic coupling between two protein positions can be experimentally verified and quantified using thermodynamic mutant cycle analysis (Vergani et al., 2005). If the two target positions indeed interact, and the strength of the interaction changes during gating, then the kinetic effects of disrupting that interaction by single mutations at either position will not be additive in the double mutant. Hypothesized interactions between amino acid side chains can be conveniently perturbed by changing the length or chemical nature of the participating side chains. Perturbing a backbone carbonyl group is less straightforward, but if the target position is located in a loop, than shortening the loop by a single-residue deletion might increase the distance to the partner position sufficiently to disrupt the hypothesized interaction (Simon and Csanády, 2021).

Following the above strategy we verified the effect on burst-state stability of disrupting the hypothesized G576-Q1371 H-bond in the human E1371Q CFTR background construct (Fig. 2E). To perturb position 1371, we shortened the side chain by substituting the glutamine with a serine (E1371S). To perturb the position of the opposing backbone carbonyl group, we shortened the D-loop by deleting residue G576 (G576Δ). We first addressed the effects of these perturbations on non-hydrolytic burst duration in macroscopic current relaxation experiments (Fig. 3A). Whereas in the E1371Q background deletion of residue G576 decreased τ_b_ by ∼20-fold (Fig. 3A, *orange* vs. *black trace*; Fig. 3B, *orange* vs. *black symbols*), in the E1371S background the effect on τ_b_ of the same deletion was only ∼2-fold (Fig. 3A, *green* vs. *blue trace*; Fig. 3B, *green* vs. *blue symbols*). From an energetic point of view, the mutation-induced shortening of τ_b_ reflects destabilization of the B_1_ state relative to the transition state for non-hydrolytic closure (T^‡^). The resulting reduction in the free enthalpy barrier for non-hydrolytic closure (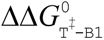; Fig. 3C, *numbers on arrows*) is quantified by the logarithm of the fractional change in τ_b_ (see Materials and Methods). The interaction energy (ΔΔ*G*_int_(B_1_→T^‡^); Fig. 3C, *purple*) is defined as the difference between 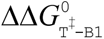 values caused by perturbation of one target position depending on whether the other target position is mutated or intact (Fig. 3C, *numbers on parallel arrows*), and quantifies the change in the strength of the G576-Q1371 interaction in the E1371Q background construct while the channel proceeds from the B_1_ state to state T^‡^. That ΔΔ*G*_int_(B_1_→T^‡^) is significantly different from zero (p=4.6·10^-5^), its magnitude (2.3±0.3 *kT*) is comparable to that of an H-bond, and its positive signature reports that the bond is present in the B_1_ state, but not in state T^‡^. Because in the IB state separation of the NBD interfaces of site 2 (Vergani et al., 2005) render formation of the inter-NBD H-bond unlikely, these results suggest that the bond is present selectively in the B state.

**Fig. 3.**
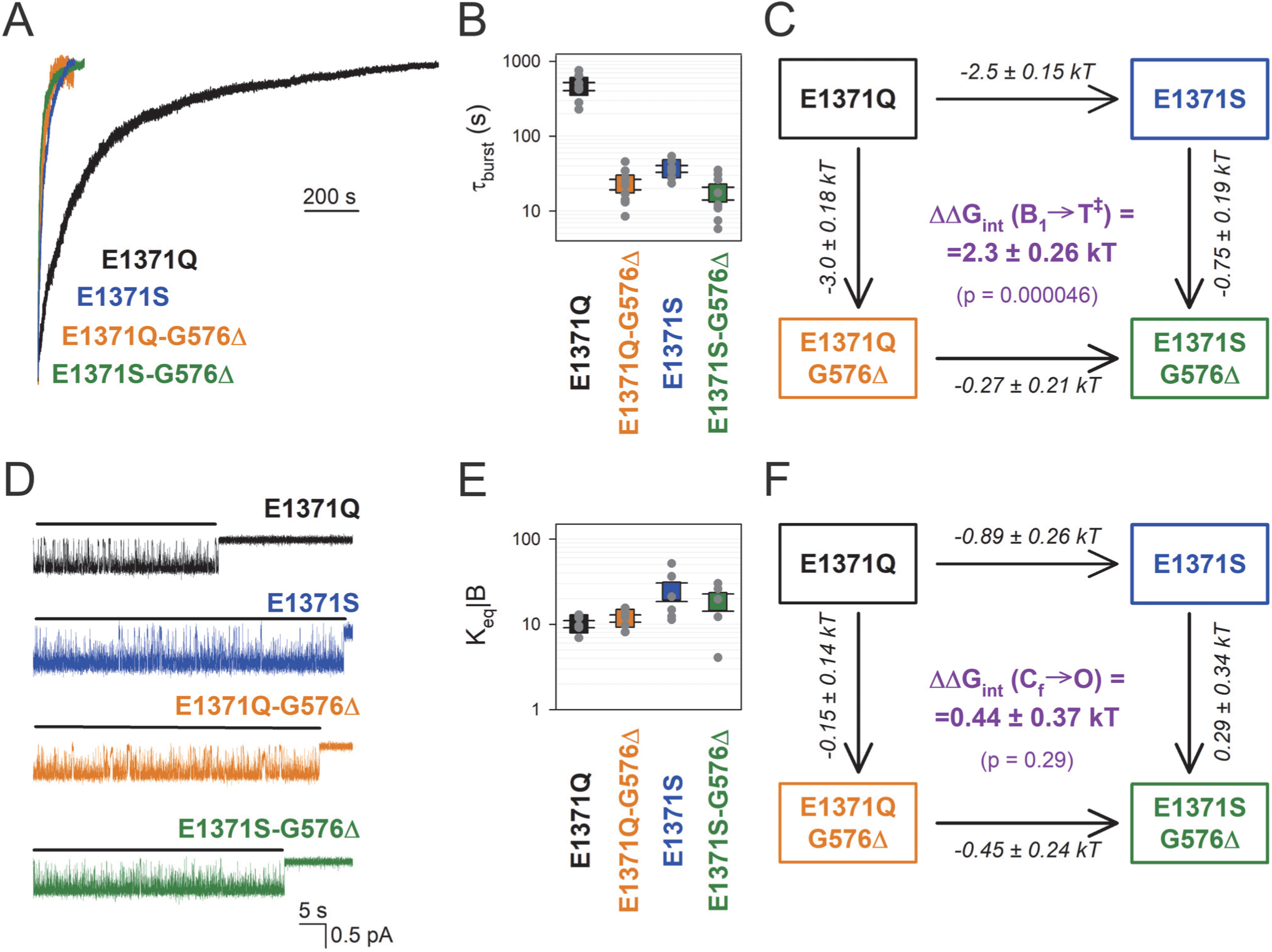
Strong coupling between positions 576 and 1371 of hCFTR E1371Q in the bursting state. *A*, Macroscopic current relaxations following ATP removal for indicated human CFTR channel mutants (*color coded*). Inside-out patch currents were activated by exposure of pre-phosphorylated channels to 5 mM ATP. Current amplitudes are shown rescaled by their steady-state values. V_m_ was -20 to -80 mV. *B*, Relaxation time constants of the currents in *A*, obtained by fits to single exponentials. Data are shown as mean±S.E.M. from 7-9 experiments using a logarithmic ordinate. *C*, Thermodynamic mutant cycle showing mutation-induced changes in the height of the free enthalpy barrier for the B_1_→IB_1_ transition (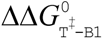, *numbers on arrows*; *k*, Boltzmann’s constant; *T*, absolute temperature). Each corner is represented by the mutations introduced into positions Q1371 and G576 of E1371Q-CFTR. ΔΔ*G*_int_(B_1_→T^‡^) (*purple number*) is obtained as the difference between 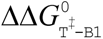 values along two parallel sides of the cycle. *D*, Currents of last open channels after ATP removal for the indicated human CFTR constructs (*color coded*). Recordings were done as in panel *A*, but on patches with smaller numbers of channels. Membrane potential was -80 mV. *Black lines* indicate the analyzed segments. *E*, Intraburst equilibrium constants obtained by dwell-time analysis for the four human CFTR constructs, plotted on a logarithmic scale. Data are shown as mean ± SEM from 5-6 experiments. *F*, Thermodynamic mutant cycle showing mutation-induced changes in the stability of the O state relative to the C_f_ state (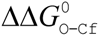, numbers on arrows; *k*, Boltzmann’s constant; *T*, absolute temperature). Each corner of the cycle is represented by the mutations introduced into positions Q1371 and G576 of E1371Q-CFTR. ΔΔG_int_(C_f_→O) (*purple number*) is obtained as the difference between 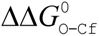 values along two parallel sides of the cycle.

### In E1371Q-hCFTR the Q1371-G576 H-bond is maintained in the flickery closed state

The bursting state is a composite state which comprises the open (O) and the flickery closed (C_f_) state. The equilibrium between those two states is reflected by the fraction of time the pore spends open within a burst (P_o|B_), which is related to the intraburst equilibrium constant through the equation *K*_eq|B_=P_o|B_/(1-P_o|B_). Intraburst gating of non-hydrolytic CFTR mutants can be conveniently studied in inside-out patches that contain small numbers of channels, through dwell-time analysis of segments of record following ATP removal in which all but one channel have terminally closed. Due to the absence of ATP such "last-channel" time windows (Fig. 3D, *black bars*) are devoid of interburst events, and allow selective collection of large numbers of open and flickery closed events (e.g., (Simon and Csanády, 2021)).

To address a potential change in the strength of the G576-Q1371 H-bond between the open and the flickery closed state of E1371Q CFTR, we compared intraburst gating of E1371Q and of the single and double target-site mutants (Fig. 3D). In contrast to the large effects of the mutations on τ_b_, the high intraburst P_o_ of the E1371Q background construct (P_o|B_=0.91±0.1 (n=5)) was not reduced in any of the single or double mutants, for all of which *K*_eq|B_ values remained within ∼2-fold (Fig. 3E). Correspondingly, a mutant cycle built on *K*_eq|B_ (Fig. 3F) revealed an interaction energy ΔΔ*G*_int_(C_f_→O) of 0.44±0.36 *kT* (Fig. 3F, *purple*), which is not significantly different from zero (p=0.29) suggesting that the strength of the interaction does not change significantly between the flickery closed and the open state.

### Catalytic glutamate mutation E1371S and Walker A mutation K1250A both completely abolish ATP hydrolysis

In WT CFTR channels the side chain of E1371 acts as the general base that deprotonates the attacking water molecule during the ATP hydrolysis reaction, whereas the side chain of the NBD2 Walker A lysine K1250 stabilizes the phosphate backbone of ATP bound in site 2, by forming strong ionic interactions with the β and γ phosphates (Fig. 4A). The K1250A and the E1371S mutations both reduce channel closing rate by ∼100-fold compared to WT CFTR, indicating that they both reduce the rate of ATP hydrolysis (*k*_1_) by at least 100-fold. If both single mutations failed to completely abolish *k*_1_ then a further slowing in *k*_1_, i.e., a further prolongation of τ_b_, would be expected in the double mutant K1250A/E1371S. However, introducing the E1371S mutation into the K1250A background did not significantly alter τ_b_ (p=0.13) (Fig. 4B, *blue* vs. *gray trace*, Fig. 4C, *blue* vs. *gray symbol*). Similarly, introduction of mutation K1250A into the E1371S background failed to prolong τ_b_. These results together indicate that ATP hydrolysis is already completely abolished in both single mutants (Fig. 4B-C, *blue* vs. Fig. 3A-B, *blue*).

**Fig. 4.**
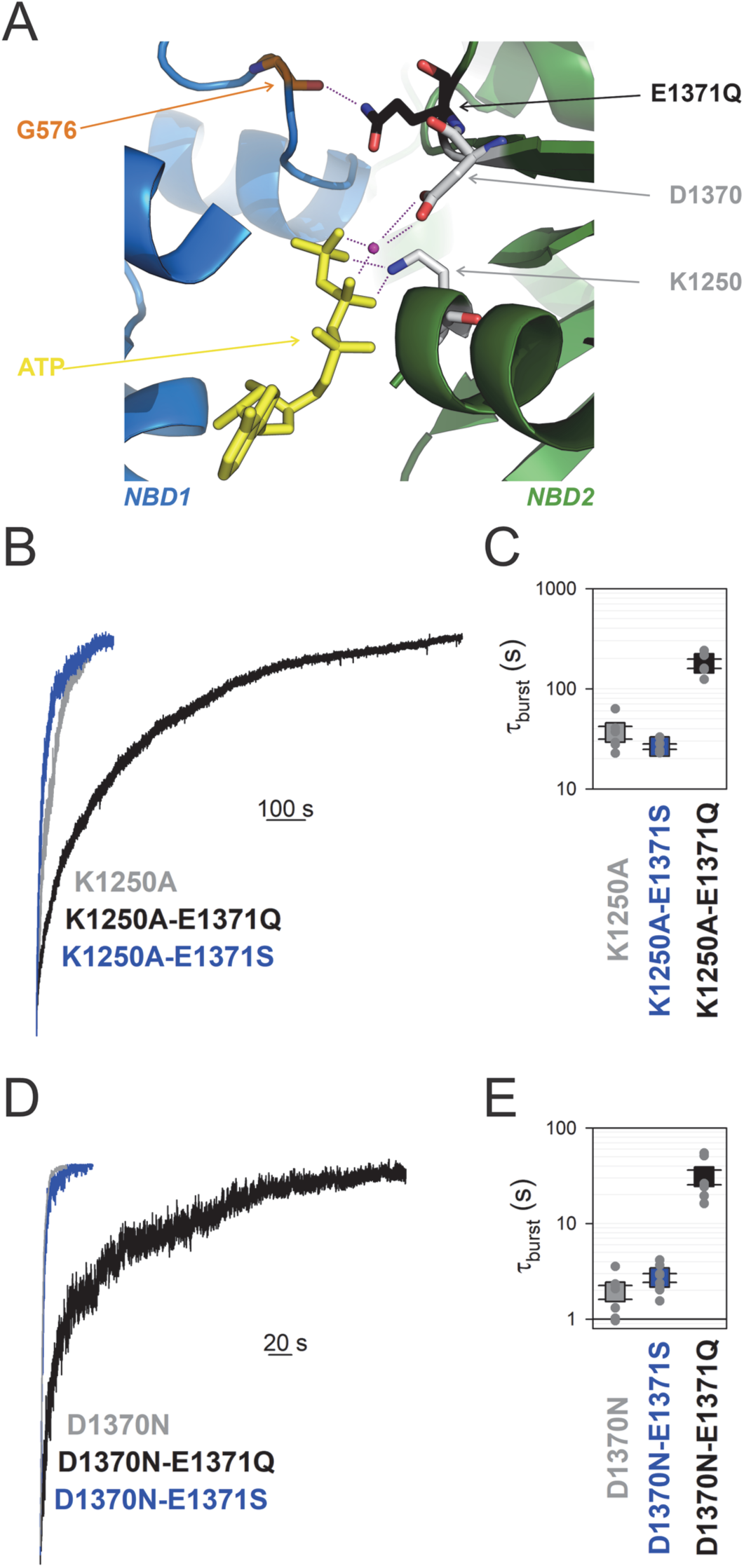
Mutation E1371Q strongly stabilizes, whereas D1370N destabilizes, bursts in other non-hydrolytic backgrounds. *A*, Close-up view of the site 2 interface in the OF structure of CFTRE1371Q (PDBID: 6msm) highlighting (in sticks) NBD1 D-loop residue G576 (*orange*), and NBD2 residues Q1371 (*black*), D1370 (*gray*), and K1250 (*gray*). ATP, *yellow sticks*; Mg2+, *purple sphere*. *B*, *D*, Macroscopic current relaxations following ATP removal for indicated CFTR channel mutants (*color coded*). Experiments were performed as in Fig. 3A, and current amplitudes are shown rescaled by their steady-state values. *C*, *E*, Relaxation time constants of the currents in *B*, *D*, obtained by fits to single exponentials, displayed on a logarithmic ordinate. Data show mean±S.E.M. from 5-8 experiments.

### Walker B mutant D1370N completely abolishes ATP hydrolysis

The conserved NBD2 Walker B aspartate D1370 coordinates the Mg^2+^ ion in site 2, required for ATP hydrolysis in all ATPases (Fig. 4A). Although in other ABC proteins D-to-N mutations at the analogous position were shown to abrogate ATP hydrolysis (Urbatsch et al., 1998; Hrycyna et al., 1999; Rai et al., 2006), for the CFTR NBD2 Walker B mutant D1370N τ_b_ is only ∼2s, i.e., ∼15 times shorter than that of most other non-hydrolytic mutants. If that relatively fast closing rate of D1370N CFTR simply reflected residual ATP-ase activity, i.e., a rate *k*_1_ of ∼0.5 s^-1^, then introducing the E1371S mutation into the D1370N background would be expected to abolish that residual activity, and prolong τ_b_ to a value near that of single mutant E1371S. However, introducing the E1371S mutation into the D1370N background did not significantly alter τ_b_ (p=0.09) (Fig. 4D, *blue* vs. *gray trace*, Fig. 4E, *blue* vs. *gray symbol*), suggesting complete lack of ATP hydrolysis in the D1370N single mutant.

### The E1371Q mutation stabilizes the bursting state of all non-hydrolytic CFTR mutants

If the E1371Q mutation indeed introduces an artificial H-bond which selectively forms in the bursting state, then introduction of that mutation into any other non-hydrolytic background construct is expected to prolong τ_b_. To verify that prediction, we introduced mutation E1371Q into both Walker mutant backgrounds. Indeed, introducing mutation E1371Q into a K1250A background further increased τ_b_ by ∼5-fold (p=2.4·10^-5^) indicating strong stabilization of state B_1_ by the E1371Q mutation (Fig. 4B, *black* vs. *gray trace*, Fig. 4C, *black* vs. *gray symbol*). Similarly, introducing the E1371Q mutation into a D1370N background further increased τ_b_ by ∼16-fold (p=1.7·10^-4^), again indicating strong stabilization of the B_1_ state (Fig. 4D, *black* vs. *gray trace*, Fig. 4E, *black* vs. *gray symbol*).

## Discussion

In ABC proteins the E-to-Q mutation of the catalytic base, a conserved glutamate that follows the Walker B aspartate, abrogates ATP hydrolysis and traps isolated NBD proteins in stable dimer form (Moody et al., 2002; Smith et al., 2002; Janas et al., 2003). For that reason, E-to-Q mutants have been adopted as model systems in a multitude of structural studies, in an attempt to trap the protein in a prehydrolytic state. Indeed, under cryo-EM conditions in the presence of ATP full-length E-to-Q mutant ABC proteins yield OF structures in which ATP is seen intact, confirming lack of hydrolysis (Liu et al., 2017; Johnson and Chen, 2018; Kim and Chen, 2018; Olsen et al., 2020; Song et al., 2021; Huang et al., 2023). In CFTR a first hint for a possible additional role, apart from simply disrupting ATP hydrolysis, of the engineered Q1371 side chain came from a recent study which compared non-hydrolytic closing rates of CFTR orthologs (Simon and Csanády, 2023). Intriguingly, whereas for hCFTR τ_b_ is >10-fold longer for the E-to-Q compared to the E-to-S mutant (Fig. 2A-B, *black* vs. *dark blue*), for the zebrafish ortholog τ_b_ of the E-to-S and E-to-Q mutants is similar (Fig. 2A-B, *gray* vs. *light blue*). Careful comparison of the dimer interface structures of OF hCFTR and zCFTR suggests the presence of a non-native H-bond in the human structure which forms between the side chain amide nitrogen of hQ1371 and the backbone carbonyl group of hG576, located at a distance of ∼3 Å in the opposing D-loop of NBD1 (Fig. 2E). Such a bond cannot form in WT CFTR, as the native glutamate side chain cannot act as a H-donor. Mutant cycle analysis indeed confirmed strong energetic coupling between positions 1371 and 576 in hCFTR/E1371Q throughout the bursting state (Fig. 3A-C), maintained even during flickery closures (Fig. 3D-F). Interestingly, formation of that H-bond in the E-to-Q mutant seems a unique feature of hCFTR, since the analogous positions are further apart in OF structures of both zCFTR (4.3Å) and of other ABCC proteins (e.g., 4.7Å in bovine MRP1, PDBID: 6bhu).

In several ABC proteins Walker A K-to-A and Walker B D-to-N mutations were shown to decrease the rate of ATP hydrolysis below the limit of detection (Ramjeesingh et al., 1999; Urbatsch et al., 1998; Hrycyna et al., 1999). However, the sensitivities of such enzymatic assays are limited, and for CFTR ATPase activity of the Walker B mutant D1370N has not been directly measured. Thus, for CFTR the >10-fold faster closing rate of K1250A and ∼200-fold faster closing rate of D1370N compared to E1371Q raised the possibility of residual ATP hydrolysis in the Walker mutants. Here we evaluated that possibility using combinations of catalytic site mutations. We found that mutation E1371S does not significantly slow closing rate of either Walker mutant (Fig. 4B-C, Fig. 4D-E), confirming that all three mutations (K1250A, D1370N, and E1371S) individually abolish ATP hydrolysis. Furthermore, the mutual lack of effect of mutation E1371S on τ_b_ of K1250A (Fig. 4B-C, *blue* vs. *gray*) and of mutation K1250A on τ_b_ of E1371S CFTR (Fig. 4B-C, *blue* vs. Fig. 3A-B, *blue*) suggests that neither of those two mutations significantly alters the stability of the B_1_ state. Thus, τ_b_ of K1250A or E1371S CFTR can be taken as relatively unbiased estimates of the life time of the B_1_ state in WT CFTR, suggesting a true *k*_-1_ of ∼0.03 s^-1^ (Fig. 5, *red*). Consistent with that notion, mutation E1371S also failed to significantly alter τ_b_ of another non-hydrolytic background construct, D1370N CFTR (Fig. 4D-E, *blue* vs. *gray*).

**Fig. 5.**
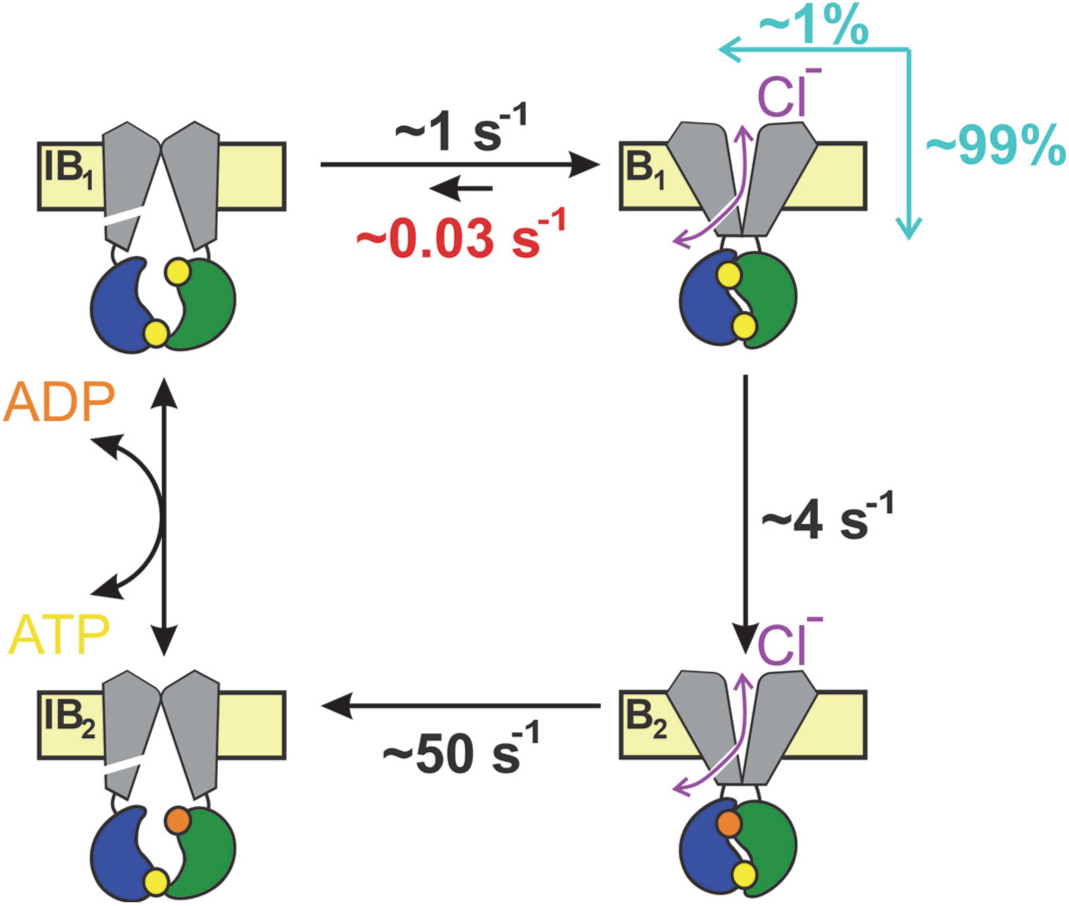
Coupling between bursts and ATP hydrolysis in WT CFTR. Quantitative CFTR gating cycle, color coding as in Fig. 1C. Numbers represent estimated microscopic rate constants for prephosphorylated WT CFTR channels gating in ATP at 25°C, with *k*_-1_ (*red*) established in the present study. *Cyan arrows* and *numbers* illustrate the fractions of bursts that are terminated by ATP hydrolysis and non-hydrolytic dissociation of composite site 2, respectively.

In contrast, mutation D1370N dramatically shortened τ_b_ of both E1371S (by ∼14-fold, Fig. 4D-E, *blue* vs. Fig. 3A-B, *blue*) and E1371Q (by ∼15-fold, Fig. 4D-E, *black* vs. Fig. 3A-B, *black*) CFTR, suggesting that the shorter τ_b_ of the D1370N single mutant compared to other non-hydrolytic mutants is explained by a large destabilizing effect of the D1370N mutation on B_1_-state stability, as opposed to substantial residual ATP-ase activity. That conclusion is consistent with the monotonically decaying distribution of D1370N burst durations, interpreted to reflect (near-) equilibrium gating of the mutant (Csanády et al., 2010).

Prompted by the discovery of the artificial H-bond in the E1371Q mutant we sought to dissect its functional effects on B_1_-state stability. Mutation E1371Q indeed greatly increases τ_b_ when introduced into any other non-hydrolytic background: by ∼13-fold in the E1371S background (Fig. 3A-B, *black* vs. *blue*), by ∼16-fold in the D1370N background (Fig. 4D-E, *black* vs. *gray*), and by ∼5-fold in the K1250A background (Fig. 4B-C, *black* vs. *gray*).

The bidirectionality of effects of different site-2 mutations on non-hydrolytic τ_b_ established here is in line with similarly bidirectional effects of different mutations at the site-1 interface documented in earlier studies. Indeed, whereas non-hydrolytic τ_b_ is decreased ∼5-10-fold by NBD1 Walker A mutation K464A (Powe, Jr. et al., 2002; Vergani et al., 2003; Csanády et al., 2010; Csanády et al., 2013), it is increased ∼3-fold by NBD2 signature sequence mutation H1348A (Szollosi et al., 2011; Csanády et al., 2013) or by N^6^-(2-phenylethyl)-ATP (P-ATP) bound in site 1 (Tsai et al., 2010; Csanády et al., 2013).

Earlier studies documented a clear pattern of larger effects on the rate of opening compared to non-hydrolytic closure (*k*_open_ vs. *k*_-1_, Fig. 1C) for mutations introduced into opposing sides of site 2 (positions 555 (Sorum et al., 2017) and 1246 (Sorum et al., 2015)), suggesting that the inter-NBD contacts across the site-2 interface are already established in the transition state (T^‡^) that separates states IB_1_ and B_1_. The lack of effect on *k*_-1_ of mutations K1250A and E1371S demonstrated here is in line with that suggestion. On the other hand, the ∼15-fold increase and decrease in non-hydrolytic τ_b_ caused by mutations E1371Q and D1370N, respectively, clearly indicate that those two side chains (located near the center of the NBD dimer interface between sites 1 and 2; Fig. 4 – Fig. Suppl. 1A) undergo significant rearrangements even during the T^‡^ ↔ B_1_ gating step. Alignment of the cryo-EM structures of NBD2 in the absence of nucleotide (from unphosphorylated WT apo structure 5uak), of NBD2-ATP without contact to NBD1 (from unphosphorylated WT ATP-bound structure 8fzq), and of NBD2-ATP tightly dimerized with NBD1 (from phosphorylated ATP-bound E1371Q structure 6msm) affords some speculation on the possible nature of those movements. For non-native Q1371 that movement might involve flipping of the mutated side chain, prompted by the vicinity of the G576 carbonyl oxygen established in state T^‡^. Indeed, whereas in the dimerized NBD2 structure the mutant Q1371 side chain is clearly resolved and points towards the NBD1 D-loop, in the ATP-bound but de-dimerized NBD2 structure no clear density is observable for the native E1371 side chain, suggesting that it is flexible or perhaps adopts a different rotameric conformation (Fig. 4 – Fig. Suppl. 1B). For the D1370 side chain the same two structures suggest a substantial movement towards the Mg^2+^ ion in the NBD-dimerized state, strengthening the D1370 – Mg^2+^ interaction (Fig. 4 – Fig. Suppl. 1B). That movement is not due to a side chain rearrangement: position 1370 is at the boundary between the "head" and "tail" subdomains of NBD2 which in all ABC proteins rotate towards each other upon ATP binding (Karpowich et al., 2001). Comparison of all three structures shows that in CFTR’s NBD2 that "subdomain closure" is fully completed only in the NBD-dimerized state (Fig. 4 – Fig. Suppl. 1C), reminiscent to earlier findings on the maltose transporter (Orelle et al., 2010). If that movement, which reduces the distance between the D1370 side chain carboxylate and Mg^2+^ from ∼5.0 to 3.7 Å, happened during step T^‡^ ↔ B_1_ then lack of the tightening D1370 – Mg^2+^ interaction in D1370N could explain selective destabilization of the B_1_ state relative to T^‡^ in the mutant.

In the past, assuming *k*_-1_ < 0.2 s^-1^ and *k*_1_ ∼4 s^-1^, a lower limit of ∼0.95 was provided for the coupling ratio (CR) of prephosphorylated WT CFTR gating at 25°C (Csanády et al., 2010). Our present estimate of *k*_-1_ ∼0.03 s^-1^ under the same conditions provides a more accurate value of CR∼0.99. Thus, only 1 out of a ∼100 bursts is terminated by non-hydrolytic separation of site 2 (Fig. 5). Furthermore, earlier estimates of temperature- and phosphorylation-dependence of rates *k*_1_ and *k*_-1_ allow some extrapolations to different experimental conditions. Based on the measured activation enthalpies (ΔH^‡^_B1→B2_ ∼70 kJ/mol, ΔH^‡^_B1→IB1_ ∼40 kJ/mol (Csanády et al., 2006)) *k*_1_ is ∼3-fold faster (∼12 s^-1^) and *k*_-1_ ∼1.9-fold faster (∼0.06 s^-1^) at 37°C, predicting a CR of ∼0.995. On the other hand, the presence of PKA slows both rates by ∼2-fold (Csanády et al., 2010; Vergani et al., 2003), predicting little effect on the CR.

A strong native H-bond between extracellular loops 1 and 6 of CFTR (between residues R117 and E1124) was shown to be present only in the open state, but disrupted during both interburst and flickery closures (Simon and Csanády, 2021). In contrast, we show here that the non-native Q1371-G576 H-bond is maintained during flickery closures, and disrupted only during interburst closures (Fig. 3). These findings are consistent with the notion that flickery closures of E1371Q CFTR involve local movements confined to the external ends of the TMD helices but no disruption of the tight NBD dimer, whereas interburst closures represent a global rearrangement during which site 2 of the NBD dimer opens up to allow nucleotide exchange. The precise extent of that separation is presently unknown: it is large enough to disrupt the site-2 interfacial H-bonds (Fig. 4 – Fig. Suppl. 1A) formed between the side chains of T1246 and R555 in WT (or K1250R) CFTR (Vergani et al., 2005) and between residues Q1371 and G576 in E1371Q CFTR (Figs. 2E, 3A-C), but too small to be detected by a pair of FRET sensors engineered into position 388 of NBD1 and position 1435 of NBD2 (Levring et al., 2023). Precisely gauging the extent of those motions will require a cryo-EM structure of the active IB state of phosphorylated CFTR in the presence of ATP.

## Materials and Methods

### Key resources table

| Reagent type (species) or resource | Designation | Source or reference | Identifiers | Additional information |
| --- | --- | --- | --- | --- |
| biological sample<br>(*Xenopus laevis*) | *Xenopus laevis* oocytes | European Xenopus Resource Centre | RRID: NXR_0.0080 | - |
| commercial assay or kit | HiSpeed Plasmid Midi Kit | Qiagen | 12643 |  |
| commercial assay or kit | QuickChange II Mutagenesis Kit | Agilent Technologies | 200524 |  |
| commercial assay or kit | mMESSAGE mMACHINE T7 Transcription Kit | ThermoFisher Scientific | AM1344 |  |
| chemical compound, drug | Collagenase type II | ThermoFisher Scientific | 17101-015 |  |
| chemical compound, drug | Adenosine 5'-triphosphoribose magnesium (ATP) | Sigma-Aldrich | A9187 |  |
| software, algorithm | Pclamp9 | Molecular Devices | RRID: SCR_011323 |  |

### Molecular biology

Mutations were introduced into the human CFTR/pGEMHE and zebrafish CFTR/pGEMHE coding sequences using the Agilent QuickChange II mutagenesis Kit. The entire coding sequences of all constructs were confirmed by automated sequencing (LGC Genomics GmbH). Plasmids were linearized using Nhe I HF (New England Biolabs) and transcribed *in vitro* (mMessage-mMachine T7 Kit, Agilent Technologies), and purified cRNA was stored at -80 °C.

### Functional expression of human CFTR constructs in Xenopus laevis oocytes

*Xenopus laevis* oocytes were removed from anaesthetized frogs following Institutional Animal Care and Use Committee guidelines, digested with collagenase (Gibco, Collagenase type II), and stored at 18°C in a solution containing (in mM) 82 NaCl, 2 KCl, 1 MgCl_2_, 1.8 CaCl_2_, and 5 HEPES (pH 7.5 with NaOH), supplemented with 50 μg/ml gentamycin. CFTR cRNA (1-20 ng) was injected in a fixed 50 nl volume (Drummond Nanoject II). Recordings were done 1-3 days following injection.

### Excised inside-out patch-clamp recording

Excised inside-out patch-clamp recordings were done as described (Simon and Csanády, 2023). The patch pipette solution contained (in mM): 138 NMDG, 2 MgCl_2_, 5 HEPES, pH=7.4 with HCl. The bath solution contained (in mM): 138 NMDG, 2 MgCl_2_, 5 HEPES, 0.5 EGTA, pH=7.1 with HCl. Following excision patches were placed into a flow chamber in which the continuously flowing bath solution could be exchanged with a time constant of <100 ms using electronic valves (ALA-VM8, Ala Scientific Instruments). Recordings were performed at 25°C, at membrane potentials of -20 to -80 mV. MgATP (5mM, Sigma-Aldrich) was diluted from a 400 mM aqueous stock solution. Channels were activated by ∼2 minute exposure to 300 nM bovine PKA catalytic subunit, prepared from beaf heart following the protocol of (Kaczmarek et al., 1980). Currents were amplified, low-pass filtered at 2 kHz (Axopatch 200B, Molecular Devices), digitized at 10 kHz (Digidata 1550B, Molecular Devices) and recorded to disk (Pclamp 11, Molecular Devices).

### Kinetic analysis of electrophysiological data

To obtain relaxation time constants, macroscopic current relaxations were fitted to single exponentials (Clampfit 11). For intraburst kinetic analysis of the last open channel, recordings were Gaussian filtered at 100 Hz, idealized by half-amplitude threshold crossing, and mean open times (τ_open_) and mean flickery closed times (τ_flicker_) obtained as the simple arithmetic averages of the mean open and closed dwell-time durations, respectively. The intraburst equilibrium constant (Fig. 3E) was calculated as *K*_eq|B_=τ_open_/τ_flicker_.

### Mutant cycle analysis

Changes in the strength of the G576-Q1371 interaction between various gating states of CFTR-E1371Q were quantitated by mutant cycle analysis as described (Mihályi et al., 2016). In brief, mutation-induced changes in the height of the transition-state barrier for non-hydrolytic closure (step B_1_→T^‡^) were calculated as 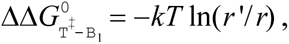 where *k* is Boltzmann’s constant, *T* is absolute temperature, and *r* and *r*’ are the rates for the B_1_→IB_1_ transition in the background construct and in the mutant, respectively. The change in the stability of the O relative to the C_f_ ground state was calculated 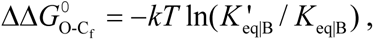 where *K*_eq|B_ and 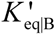 are the equilibrium constants for the C_f_↔O transition in the background construct and in the mutant, respectively. Interaction free energy (ΔΔ*G*_int_) was defined as the difference between ΔΔ*G*^0^ values along two parallel sides of a mutant cycle. All ΔΔ*G* values are given as mean±S.E.M; S.E.M. values were estimated assuming that *r* and *K*_eq|B_ are normally distributed random variables, using second-order approximations of the exact integrals (Mihályi et al., 2016).

### Statistics

All values are given as mean±S.E.M., with the numbers of independent biological replicates provided in the figure legends. Significances were evaluated using Student’s paired t-test.

## Acknowledgements

Supported by EU Horizon 2020 Research and Innovation Program grant 739593, Cystic Fibrosis Foundation Research Grant CSANAD21G0, and NKFIH KKP_22 grant 144199 to LC. M.A.S. received support from the ÚNKP-22-3-II-SE-12 New National Excellence Program of the Ministry for Innovation and Technology from the source of the National Research, Development and Innovation Fund.

## Author contributions

M.A.S. and L.C. designed the project. I.I. and A.S. purified the bovine PKA catalytic subunit. M.A.S. generated mutant constructs, performed electrophysiological experiments and analyzed the data. M.A.S. and L.C. interpreted the results and wrote the manuscript.

## Supplementary Figures

**Fig. 2 – Fig. Suppl. 1.**
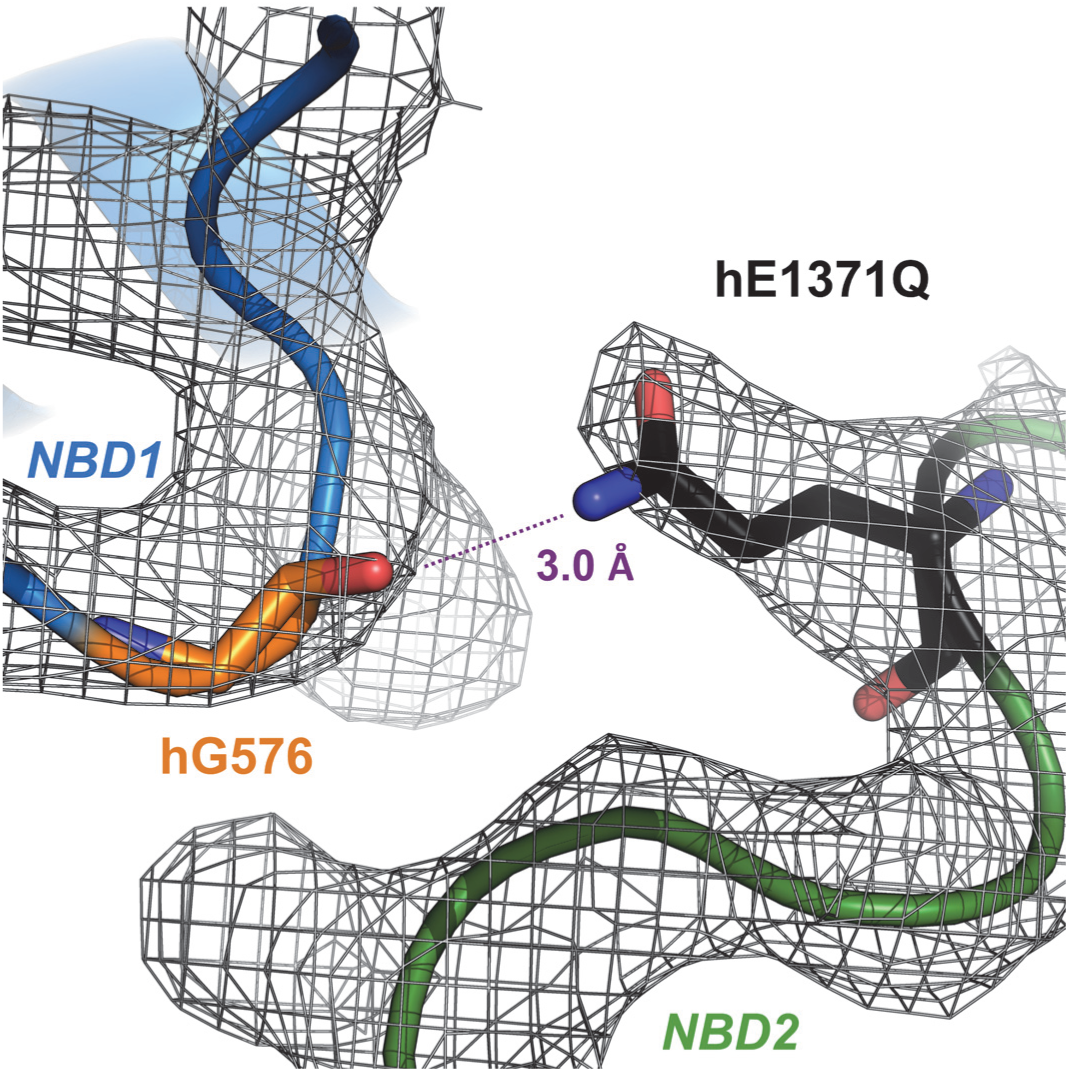
Experimental electrondensities around residues G576 and Q1371 in OF structure of hCFTR E1371Q. Electrondensities (*gray mesh*, EMD-9230) and atomic model (*sticks*, PDBID: 6msm) of residues G576 and Q1371 in OF CFTR-E1371Q.

**Fig. 4 – Fig. Suppl. 1.**
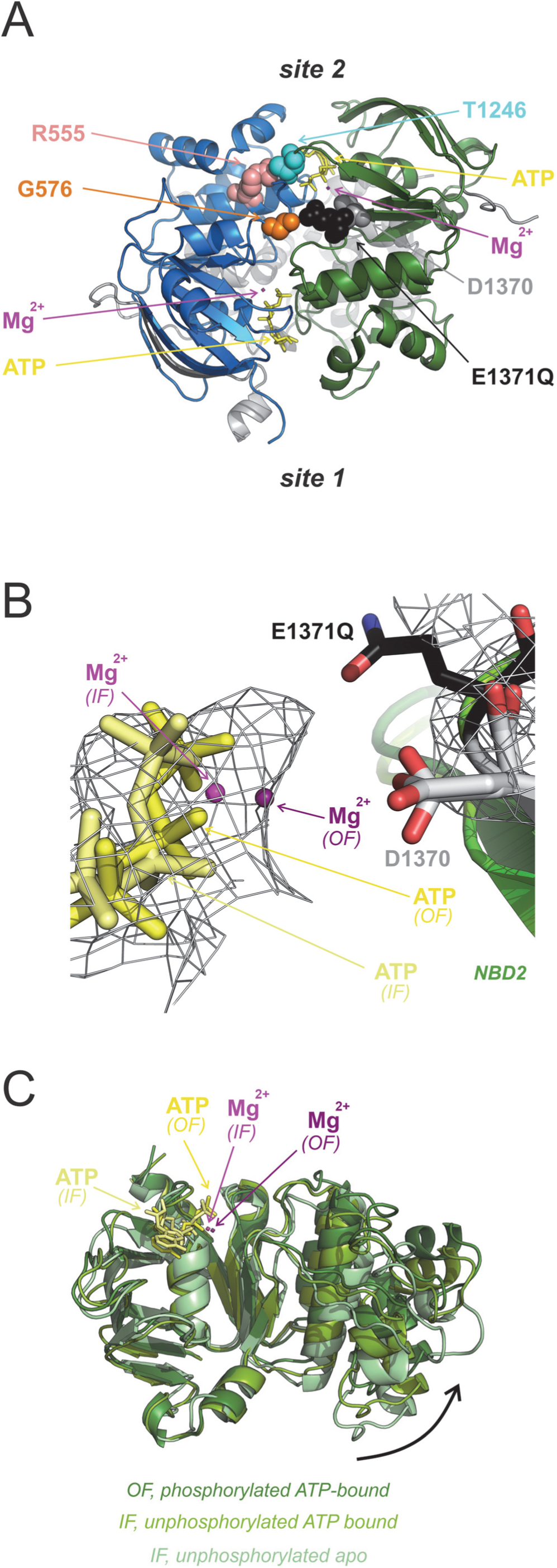
Intra-NBD2 movements associated with ATP binding and tight NBD dimerization. *A*, Location of positions 576, 1370, and 1371 (*colored spacefill*) relative to composite sites 1 and 2 in the tight NBD dimer (PDBID: 6msm). Interacting positions 555 and 1246 (Vergani et al., 2005) are also shown in colored spacefill. *B*, Atomic models (*sticks*) of ATP (*shades of yellow*), Mg^2+^ (*shades of purple*), and residue D1370 from ATP-bound non-phosphorylated IF WT CFTR (PDBID: 8fzq) and phosphorylated ATP-bound OF E1371Q CFTR (PDBID: 6msm) overlayed with the electrondensity for the IF structure (EMD-29637). The core subdomain of NBD2 from 6msm was aligned with that of 8fzq in Pymol. Residue 1371 (*black sticks*) is shown only for the OF structure. Note lack of electrondensity in the IF map in the region occupied by the Q1371 side chain in OF CFTR. *C*, NBD2 structures of non-phosphorylated apo IF CFTR (6uak; *light green*), non-phosphorylated ATP-bound IF CFTR (8fzq; *medium green*), and phosphorylated ATP-bound OF E1371Q-CFTR (6msm; *dark green*), aligned through their core subdomains. ATP (*shades of yellow*), and Mg^2+^ (*shades of purple*) of the ATP-bound structures are shown in stick representation. Note completion of subdomain closure (*curved arrow*) in the OF structure.

